# Antiviral Activity of Rosemary Extract Against Zika Virus in Human Dermal Fibroblasts

**DOI:** 10.1101/2025.01.30.635802

**Authors:** Jordan T. Masi, Robert W.E. Crozier, Jeremia M. Coish, Natalie J. Hicks, Carmela A. Lowes, Evangelia Tsiani, Dustin Duncan, Edward I. Patterson, Adam J. MacNeil

## Abstract

Flaviviruses have increasingly emerged and re-emerged in recent decades, infecting millions of people annually. Zika virus (ZIKV) is particularly concerning due to its associated pathological complications, including microcephaly in newborns and Guillain-Barré syndrome in adults, posing a significant threat to public health. Despite efforts made by the scientific community, no licensed drugs against flaviviruses have been developed. Medicinal plants show promise as a novel source of antiviral agents, as they possess a diverse array of biologically active secondary metabolites, making them potential candidates for therapeutic use. Here, we sought to investigate the antiviral potential of rosemary extract (RE) against ZIKV in human dermal fibroblasts (HFF-1), one of the earliest cellular targets of infection. ZIKV was treated with various concentrations of RE or its major polyphenols, including rosmarinic acid (RA), carnosic acid (CA), and carnosol (CO), and the infectivity of each sample was measured by plaque reduction assay. To evaluate the impact of RE on different stages of the ZIKV replication cycle, HFF-1 cells were treated before, during, and after infection, or the virus was treated before infection. RE exerted potent virucidal activity against ZIKV in both Vero and HFF-1 cells by directly acting on virus particles before infection. Importantly, RE significantly inhibited the later stages of the virus replication cycle by interfering with post-entry mechanisms within the host cell. Moreover, major RE-derived polyphenols CA and CO, but not RA, were shown to significantly reduce infectivity when ZIKV was pre-treated with each compound individually. Overall, RE significantly impairs ZIKV infection *in vitro* by directly interacting with virus particles prior to adsorption and interfering with post-entry processes of the viral replication cycle. This study highlights the antiviral potential of RE and its individual components, wFigurearranting further investigation into the mechanisms underlying their activity and their effects on other medically important flaviviruses.

## Introduction

Zika virus (ZIKV) is a mosquito-borne virus that has received significant attention in recent years. ZIKV belongs to the *Flavivirus* genus within the *Flaviviridae* family, which includes dengue virus (DENV), yellow fever virus (YFV), Japanese encephalitis virus (JEV), West Nile virus (WNV), and other well-known human pathogens^1^. ZIKV was originally isolated from a Rhesus macaque monkey in Uganda when studying the ecology of YFV in 1947. After its discovery, ZIKV remained relatively unknown, sporadically causing minor outbreaks and mild febrile illness in few individuals^2,3^. The first major outbreak of ZIKV occurred on Yap Island in 2007, where close to 75% of the population was infected^4^. This was followed by an outbreak in French Polynesia from 2013 to 2014 and the largest recorded outbreak in Brazil in 2015, where approximately 1.3 million people were infected^3,5^. These outbreaks were linked to a significant increase in microcephaly in neonates of infected mothers and Guillain-Barré syndrome in adults, sparking a global panic^6^. Due to its rapid spread and associated neurological complications, ZIKV was declared a Public Health Emergency of International Concern by the World Health Organization (WHO) in February 2016^7^.

ZIKV is a positive-sense single-stranded RNA virus with a genome of approximately 11 kb. The genome encodes a single polyprotein that is proteolytically cleaved into three structural proteins, including envelope, pre-membrane, and capsid proteins, along with seven non-structural (NS) proteins, including NS1, NS2A, NS2B, NS3, NS4A, NS4B, and NS5^3,4^. ZIKV is primarily transmitted by female *Aedes aegypti* mosquitoes, which inject the virus into the skin of the host during a blood meal^8^. Throughout this process, ZIKV encounters cells in the epidermal and dermal layers of the skin, including keratinocytes, fibroblasts, Langerhans cells, dendritic cells, and mast cells, representing the first line of defense against the virus^9,10^. Fibroblasts are the most abundant cell type found in the dermis and are responsible for producing and maintaining the extracellular matrix^11^. It has been well characterized that fibroblasts are highly permissive to multiple strains of ZIKV and serve as a primary target for viral replication at the site of inoculation. Moreover, multiple pro-inflammatory genes are upregulated in fibroblasts infected with ZIKV, suggesting they contribute to the local inflammatory response in the early stages of infection^10,12^. Fibroblasts are thus a compelling *in vitro* model for studying potential antiviral agents against ZIKV.

As of May 2024, 92 countries and territories throughout five of the six WHO regions have recorded autochthonous mosquito-borne transmission of ZIKV. Additionally, competent populations of *Aedes aegypti* mosquitoes have been established in 60 countries and territories globally, raising concerns that autochthonous transmission may be introduced in these areas^3,12^. Despite an extensive worldwide research effort, there are currently no vaccines approved to prevent ZIKV infection and no specific antivirals available for any flavivirus, leaving supportive care as the only option to manage symptoms. Given the severe complications associated with ZIKV infection and the ongoing spread of flaviviruses to new regions, there is an urgent need to develop effective antiviral agents to control future outbreaks^13–15^.

An attractive approach to developing therapeutics is the use of medicinal plants, as they possess a diverse array of bioactive secondary metabolites^14^. These compounds have been shaped by millions of years of evolution to provide plants selective advantages, including protection against oxidative stress, ultraviolet radiation, predators, and infection^16,17^. Historically, humans have used plants as a source of medicines for treating various diseases, a practice that continues today^18^. Plants have played a significant role in modern drug discovery, making up approximately 6% of new molecular entities approved by the U.S. Food and Drug Administration between the years 2000 and 2016^18,19^. In fact, many drugs currently available on the market were isolated from plants, including the anti-inflammatory acetylsalicylic acid (aspirin), the narcotic morphine, and the anti-glaucoma drug pilocarpine^14,20^. In the context of mosquito-borne infections, quinine, derived from the bark of the cinchona tree, stands out as one of the most important anti-malarial drugs in history and is listed on the WHO’s Model List of Essential Medicines for severe malaria^21^. Beyond their well-documented antioxidant and anti-inflammatory properties, several plant-derived secondary metabolites have demonstrated antiviral activity *in vitro* and *in vivo*, including quercetin and fisetin against DENV2, baicalein against JEV and influenza virus, and epigallocatechin-3-gallate against herpes simplex virus (HSV), Epstein Barr virus, and DENV, to name a few^22,23^. However, relatively few studies have explored the antiviral potential of medicinal plants and their individual components against ZIKV, highlighting a gap in the field.

*Rosmarinus officinalis* L. (rosemary) is a medicinal herb belonging to the *Lamiaceae* family and is native to the Mediterranean region. Rosemary extract (RE) is widely used as a natural preservative in food processing and has been extensively studied for its antioxidant, anti-inflammatory, antibacterial, antitumor, neuroprotective, and other therapeutic properties^24–26^. These can be attributed to its rich polyphenolic composition, including phenolic acids like rosmarinic (RA), ursolic, and caffeic acids, as well as phenolic diterpenoids like carnosic acid (CA), carnosol (CO), and rosmanol^27^. Although the antibacterial activity of RE has been well characterized^28^, little is known about its antiviral activity. To the best of our knowledge, no studies have explored the antiviral activity of RE in the context of flavivirus infection, although it has been shown to be effective against human respiratory syncytial virus (hRSV)^29^ and HSV^30–33^. Additionally, RA, a major polyphenol in RE, was identified as a potential antiviral agent against ZIKV and DENV *in silico*^34–36^. RA has also been shown to inhibit all four serotypes of DENV *in vitro*^36^ and significantly reduce mortality, viral loads, and pro-inflammatory cytokine levels in mice infected with a lethal dose of JEV^37^. In the current study, we demonstrate that RE potently inhibits ZIKV infection by directly interacting with virus particles before infection and disrupting later stages of the viral replication cycle. Furthermore, we found that CA and CO, but not RA, exert virucidal activity against ZIKV. These findings highlight the antiviral potential of RE and its individual components, warranting further investigation into their mechanism of action and activity against other flaviviruses.

## Materials and Methods

### Cells and virus

Human foreskin fibroblast cells (HFF-1; ATTC, SCRC-1041) and Vero E6 cells (African green monkey kidney cells; ATTC, CRL-1586) were maintained in Dulbecco’s modified Eagle’s medium (DMEM; Sigma, D6546) supplemented with 10% Serum Plus II (Sigma, 14009C) and 1% penicillin/streptomycin (pen/strep; Gibco, 15140-122) at 37°C with 5% CO_2_. The PRAVBC59 strain of ZIKV (ATCC, VR1843) was propagated in Vero cells. Briefly, Vero monolayers were infected with ZIKV at a multiplicity of infection (MOI) of 0.01 and incubated at 37°C and 5% CO_2_ for 1 h. The viral inoculum was removed, and cells were incubated for 5 days in culture medium containing 2% Serum Plus II and 1% pen/strep. Viral stocks were quantified by plaque assay, aliquoted, and stored at -80°C.

### Reagent preparation

Rosemary extract was prepared by grinding and passing whole dried *Rosmarinus officinalis* L. leaves (purchased from Sobeys, Mississauga, ON, Canada; product of Turkey) through a mesh sieve. Extraction was performed following protocols established by the US National Cancer Institute. Ground leaves (5 g) were steeped overnight in 30 mL of dichloromethane-methanol (1:1). A slight vacuum was used to collect the filtrate, and a methanol extraction was performed for 30 min. The solvent was removed by rotary evaporation. For antiviral experiments, RE was dissolved in dimethyl sulfoxide (DMSO; Sigma, D2650) at a final concentration of 100 mg/mL and RA (Sigma, R4033), CA (Sigma, C0609), and CO (Sigma, C9617) were dissolved in DMSO at a final concentration of 100 mM. Aliquots were stored protected from light at -20°C until further use. For HPLC analysis, RE was dissolved in DMSO (Sigma, 34869) at 50 mg/mL and authentic standards of RA (Cayman Chemicals, 70900), CA (Cayman Chemicals, 89820), and CO (Cayman Chemicals, 89800) were dissolved in DMSO at 5 mg/mL immediately before use.

### Cell viability assay

To determine the impact of RE treatment on cell viability, HFF-1 and Vero cells were seeded in 96-well tissue culture plates and incubated at 37°C with 5% CO_2_ until ∼90% confluent. Cells were treated in triplicate with DMEM containing 20 to 200 µg/mL of RE for 1 h at 37°C with 5% CO_2_, using 1% DMSO as a vehicle control. Subsequently, the medium was replaced with fresh culture medium containing 10% Serum Plus II and 1% pen/strep, and cells were incubated for either 96 h (HFF-1) or 120 h (Vero) at 37°C with 5% CO_2_. These conditions correspond to cells treated before or during infection or infected with pre-treated ZIKV. Additionally, HFF-1 cells were incubated with 20 to 200 µg/mL of RE in DMEM with 10% Serum Plus II and 1% pen/strep for 96 h, corresponding to cells treated after infection in the time-of-addition analysis. Following incubation, culture medium was replaced with 100 µL of fresh medium and 10 µL of WST-1 cell proliferation reagent (Roche, 5015944001). After 2 h, absorbance was measured at 460 nm using a spectrophotometer (Bio-Tek, Synergy HT-1, 191356) and triplicates were averaged. The cytotoxic concentration of RE that reduced viability by 50% (CC50) was calculated by non-linear regression analysis (GraphPad Prism).

### Plaque reduction assay

Plaque reduction assay was used to screen the antiviral activity of RE and three major polyphenol constituents commonly reported in the literature^38–41^, including RA, CA, and CO. 100 µL of ZIKV (290,000 PFU) was treated with 25 to 200 µg/mL of RE or 25 to 100 µM of each compound individually for 1 h at 37°C with 5% CO_2_, using 1% DMSO as a vehicle control. The infectivity of each sample was then quantified by plaque assay. The inhibitory concentration that reduced plaque numbers by 50% (IC_50_) was calculated by non-linear regression analysis (GraphPad Prism).

### Time-of-addition analysis in HFF-1 cells

To determine which stages of infection are targeted by RE, time-of-addition analysis was performed. Briefly, HFF-1 cells were seeded on 24-well plates and incubated at 37°C with 5% CO_2_ until ∼90% confluent. Cells were infected with ZIKV at an MOI of 1 under four different treatment conditions, with 1% DMSO included as a vehicle control. 1) Cells were pre-treated with 20 to 100 µg/mL of RE for 1 h at 37°C. The supernatant was removed, and wells were washed once with 1X phosphate-buffered saline (PBS; Gibco, 10010023) before viral inoculation. 2) Cells were infected with ZIKV and simultaneously treated with 20 to 100 µg/mL of RE for 1 h at 37°C. 3) Cells were infected with ZIKV for 1 h at 37°C, the inoculum was removed, and culture medium containing 20 to 100 µg/mL of RE was added to each well for the entire post-infection period. 4) ZIKV was incubated with 20 to 100 µg/mL of RE for 1 h at 37°C before being added to the cells. In all treatment conditions, cells were washed once with PBS after the 1 h adsorption period to remove unbound ZIKV. Next, 500 µL of culture medium with 10% Serum Plus II and 1% pen/strep was added to each well and cells were incubated at 37°C and 5% CO_2_. Supernatant was collected, aliquoted, and stored at -80°C after 0 and 96 h post-infection. Viral titers were quantified by plaque assay and genome copies were quantified by two-step reverse transcription-quantitative polymerase chain reaction (RT-qPCR). The inhibitory concentration that reduced viral titers or genome copies by 50% (IC_50_) was calculated by non-linear regression analysis (GraphPad Prism).

### Plaque assay

Vero E6 cells were seeded in 12- and 24-well plates and incubated at 37°C with 5% CO_2_ until 90% confluent. Serial 1:10 dilutions were prepared for pre-treated ZIKV stocks (plaque reduction assay) or supernatant harvested from HFF-1 cells (time-of-addition analysis) 96 h post-infection, whereas a 1:2 dilution was prepared for samples collected 0 h post-infection. Cells were infected in duplicate with 200 µL (1:10 dilutions) or 100 µL (1:2 dilutions) of each dilution in 12- or 24-well plates, respectively. Following a 1 h adsorption period at 37°C, viral inoculum was removed, and cells were overlaid with 1 mL (12-well) or 0.5 mL (24-well) of a 1:1 mixture of carboxymethylcellulose (CMC) and DMEM with 2% Serum Plus II and 1% pen/strep. Plates were incubated at 37°C with 5% CO_2_ for 5 days for plaque development, the overlay was discarded, and wells were washed twice with PBS. Cells were simultaneously fixed and stained with 1% crystal violet dissolved in 8% formaldehyde and 30% ethanol for 30 min at room temperature. Excess stain was removed using tap water and plaques were counted. Plaque forming units per mL (PFU/mL) was calculated for each sample using the following equation:

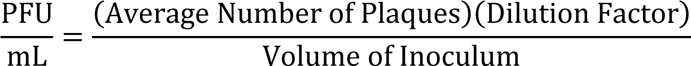

### RNA isolation and two-step RT-qPCR

Viral RNA was isolated from cell-free supernatant collected 96 h post-infection using the QIAamp Viral RNA Kit (Qiagen, 52904) according to the manufacturer’s instructions. cDNA was generated by reverse transcription using RNA to cDNA EcoDry Premix kits (Takara Bio, 639549) in a SimpliAmp Thermal Cycler (Applied Biosystems, 228001548) and diluted 1:20 in molecular grade water (Sigma, 693520). The StepOnePlus Real-Time PCR System (Applied Biosystems, 272004476) was used to perform qPCR with KAPA SYBR Fast Master Mix (KAPA Biosystems, KM4103) and amplification efficiency optimized primers at a final reaction volume of 10 µL (1 µL cDNA + 9 µL master mix). The following forward and reverse primer set was used: 5’-GCAAACGCGGTCGCAAACCT-3’ and 5’-TGCTAACGCGAAGCCAGGGT-3’ (Integrated DNA Technologies)^42^. To determine the relative number of ZIKV copies in cell-free supernatant, a standard curve was generated using ZIKV gBlock gene fragments (Integrated DNA Technologies).

### High-performance liquid chromatography (HPLC)

The polyphenolic composition of RE was analyzed using a Dionex UltiMate 3000 UHPLC system (Thermo Scientific) equipped with a Kinetex C18 column (100 mm × 4.6 mm × 2.6 µm; Phenomenex, 00D-4462-E0) maintained at 30°C. The mobile phase consisted of (A) water (Sigma, 270733) containing 1% (v/v) glacial acetic acid (Sigma, AX0074-6) and (B) acetonitrile (VWR, BDH83639.400) containing 1% (v/v) glacial acetic acid. The following gradient elution was performed: 0–25 min, 95–5% A; 25–28 min, 5% A; 28–30 min, 5–95% A; 30–35 min, 95% A. RE was dissolved in DMSO at 50 mg/mL and diluted ten-fold in 50:50 (v:v) A:B to a final concentration of 5 mg/mL for analysis. Authentic standards of RA, CA, and CO were prepared in DMSO at 5 mg/mL and diluted ten-fold in 50:50 (v:v) A:B to 500 µg/mL, followed by serial two-fold dilutions. The injection volume was 5 µL, with a flow rate of 1.0 mL/min and detection wavelength of 280 nm. Peaks were identified by comparing retention times with those of authentic standards. Quantification was performed using the external standardization method in Chromeleon (v.7.1.1.1127), with standard curves generated by plotting each concentration (31.75, 62.5, 125, 250, and 500 µg/mL) against the relative peak area.

### Statistical analysis

One-way ANOVA analysis was used to evaluate WST-1 viability assay and RT-qPCR data, and two-way ANOVA analysis was used to evaluate plaque assay data. Multiple comparisons were performed using Dunnett’s *post-hoc* test where appropriate. CC50 and IC_50_ values were determined by analyzing normalized data (% control) using non-linear regression (variable slope). Results from all experiments were expressed as mean ± SEM and considered significant if p<0.05 between the vehicle control (DMSO) and RE or polyphenol treated groups. Statistical analyses were conducted using GraphPad Prism (v.10.5.0).

## Results

### Cytotoxicity of RE in cell culture

WST-1 assay was used to evaluate the cytotoxic effects of RE treatment on HFF-1 and Vero cells. Both cell lines were exposed to increasing concentrations of RE (20–200 µg/mL) or an equivalent volume of DMSO (vehicle control). Incubation periods were selected to reflect the conditions used in the different functional assays. Results are expressed as a percentage of the vehicle control (set to 100% viability) for each independent experiment. No difference in viability was observed in HFF-1 cells treated with 20 to 80 µg/mL of RE for 1 h followed by a 96 h incubation compared to DMSO treated cells. However, there was a significant decrease in viability beginning at 100 µg/mL, with a CC50 of 120.2 µg/mL (Fig. 1a and 1c). Comparatively, HFF-1 cells treated for the entire 96 h incubation period showed a slight increase in cell viability at 40 µg/mL (p<0.01). There was a significant decrease in viability beginning at 80 µg/mL, with a CC50 of 107.3 µg/mL (Fig. 1b and 1d). Vero cells were unaffected by treatment with 20 µg/mL of RE for 1 h followed by a 120 h incubation (Fig. 2a).

**Figure 1:**
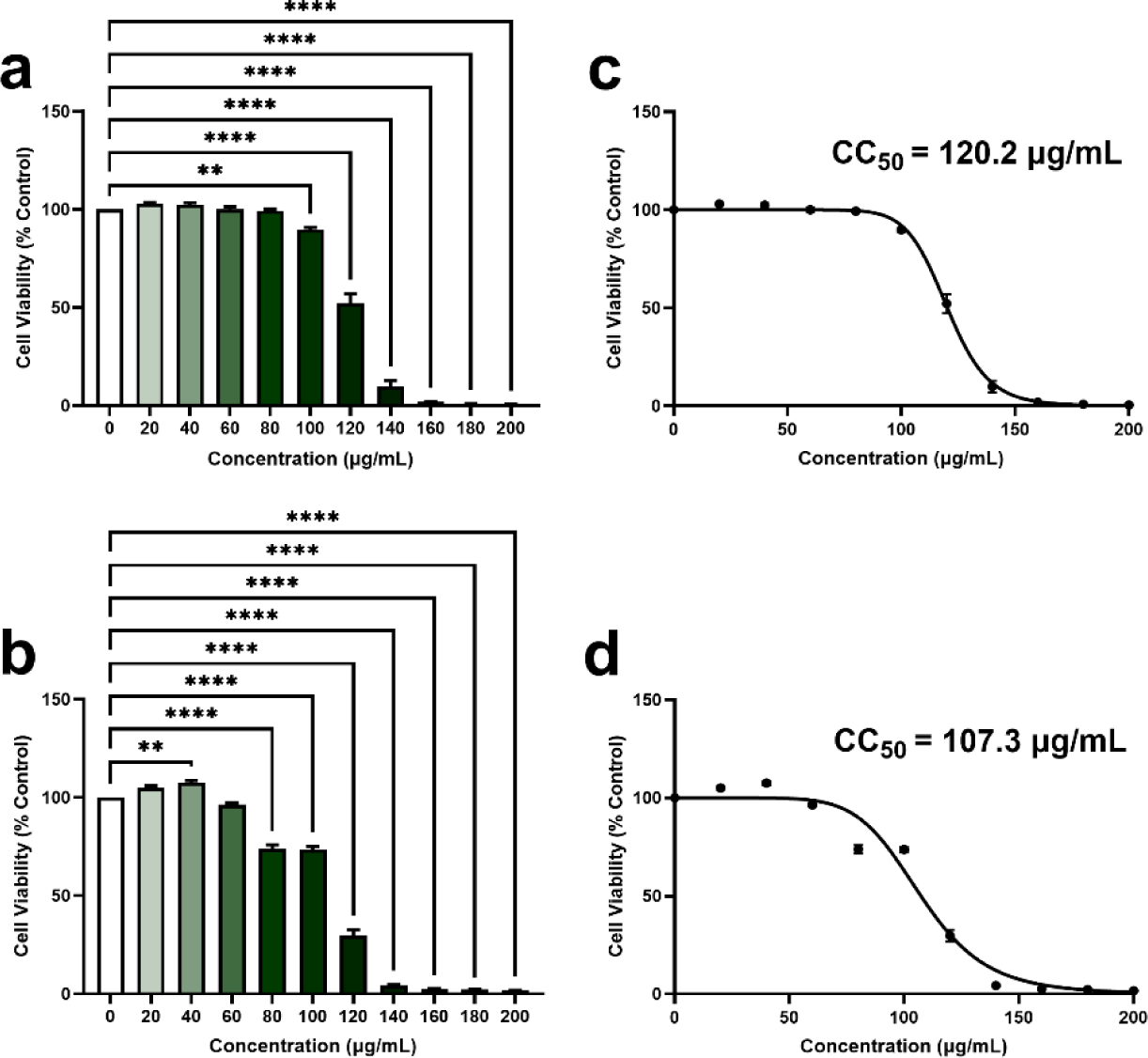
Cell viability of HFF-1 cells treated with increasing concentrations of RE. (a) HFF-1 cells were incubated with increasing concentrations of RE or a vehicle control (DMSO) for 1 h followed by 96 h with fresh medium. (b) HFF-1 cells were incubated with increasing concentrations of RE or a vehicle control (DMSO) for 96 h. Cell viability was determined by a WST-1 assay. Data are expressed as mean % of control ± SEM for n = 3 independent experiments. A one-way ANOVA and Dunnett’s multiple comparisons test were used to determine differences in cell viability following RE incubation. **p<0.01, ****p<0.0001 relative to HFF-1 cells incubated with DMSO (0 µg/mL). CC50 values for HFF-1 cells treated for (c) 1 h or (d) 96 h were calculated by non-linear regression analysis (inhibitor vs normalized response – variable slope).

**Figure 2:**
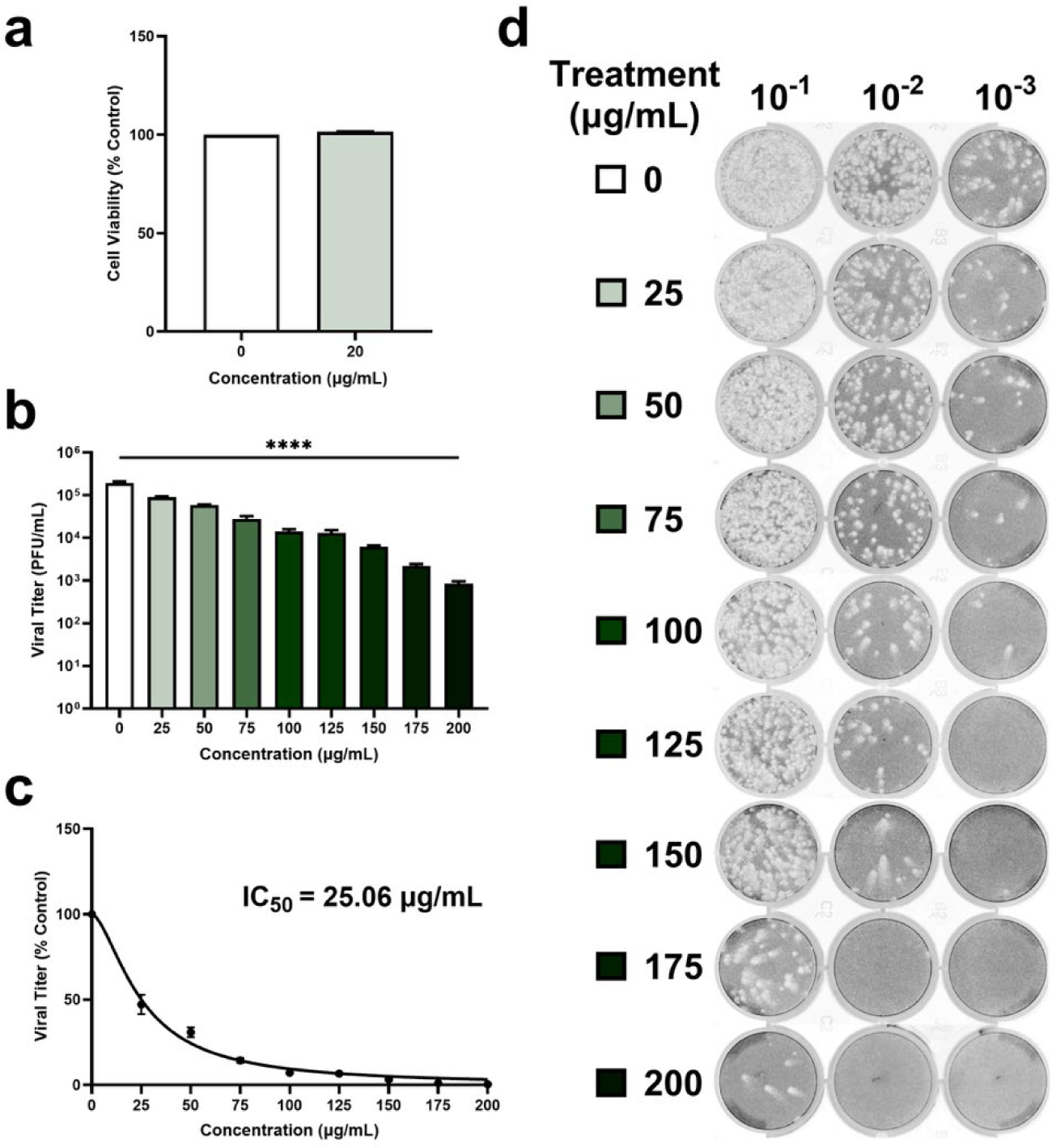
RE inhibits ZIKV replication in Vero cells in a dose-dependent manner. (a) Vero cells were incubated with increasing concentrations of RE or a vehicle control (DMSO) for 1 h followed by 120 h with fresh medium. Cell viability was determined by a WST-1 assay. Data are expressed as % of control ± SEM for n = 3 independent experiments. Vero cells were exposed to a maximum concentration of 20 µg/mL in the plaque reduction assay; thus, the corresponding representative data is shown. A one-way ANOVA and Dunnett’s multiple comparisons test were used to determine differences in cell viability following RE incubation. (b) As an initial screen, the virucidal activity of RE was evaluated by plaque reduction assay. ZIKV (290,000 PFU) was incubated with increasing concentrations of RE or an equal volume of a vehicle control (DMSO) for 1 h. The viral titer of each sample was then determined by plaque assay. Data are expressed as mean PFU/mL ± SEM for n = 3 independent experiments, each performed in duplicate. A one-way ANOVA and Dunnett’s multiple comparisons test were used to determine differences in viral titers following RE treatment. ****p<0.0001 relative to ZIKV incubated with DMSO (0 µg/mL) for all concentrations tested. (c) The IC_50_ value was calculated by non-linear regression analysis (inhibitor vs normalized response – variable slope). (d) Plaque assay images were taken using an Amersham Typhoon Biomolecular Imager and represent wells from one independent experiment in its entirety.

### RE dose-dependently inhibits ZIKV replication in Vero cells

To test the potential antiviral activity of RE, ZIKV was pre-treated with increasing concentrations of RE (25–200 µg/mL) or an equivalent volume of DMSO for 1 h. The viral titer of each sample was then quantified by plaque assay. Results demonstrated a significant dose-dependent decrease in infectious virions relative to the control (Fig. 2b, p<0.0001 for all tested concentrations). Concentrations of 25, 50, 75, 100, 125, 150, 175, and 200 µg/mL caused a 52.77 ± 5.66%, 69.06 ± 2.88%, 85.63 ± 1.49%, 92.83 ± 0.51%, 93.28 ± 0.63%, 96.74 ± 0.29%, 98.86 ± 0.06%, and 99.55 ± 0.10% reduction in ZIKV titers, respectively (Fig. 2c, IC_50_ = 25.06 µg/mL). Due to the nature of this assay, the selectivity index (SI) was not calculated. The highest concentration of RE that ZIKV was pre-treated with was 200 µg/mL, but this was serially diluted ten-fold before being added to Vero cells. As a result, Vero cells were exposed to a maximum concentration of 20 µg/mL of RE, which was not cytotoxic (Fig. 2a).

### RE directly interacts with virus particles and inhibits post-entry mechanisms in fibroblasts

To determine the antiviral potential of RE against ZIKV infection and its effect on different stages of the viral replication cycle, HFF-1 cells were treated with RE before, concurrent with, or after viral inoculation, or ZIKV was pre-treated with RE prior to infection. Increasing concentrations of RE ranging from 20 to 100 µg/mL or an equivalent volume of DMSO were used in all experiments. Based on what is frequently used in the literature^10,43–46^, cells were infected at an MOI of 1 and cell-free supernatant was collected 0 and 96 h post-infection, which is when ZIKV titers peak in HFF-1 cells (Supplemental Fig. 1). The collected supernatants were aliquoted and used for both plaque assay and RT-qPCR analysis. Percent reductions were calculated relative to the DMSO control group within each respective treatment condition. There were no statistically significant differences in viral titers between samples collected 0 h post-infection for all treatment conditions (data not shown). No antiviral effects were observed in HFF-1 cells treated with RE for 1 h prior to infection (Fig. 3a and 3b). Treatment of HFF-1 cells during the adsorption period significantly enhanced ZIKV replication, with viral titers increasing by 195.47 ± 53.63% (p<0.0001), 104.50 ± 18.36% (p<0.01), 98.68 ± 27.32% (p<0.01), 195.89 ± 65.67% (p<0.0001), and 88.65 ± 41.33% (p<0.05) and genome copy numbers increasing by 320.97 ± 36.65% (p<0.001), 274.66 ± 11.34% (p<0.01), 247.84 ± 61.02% (p<0.01), 303.73 ± 79.16% (p<0.01), and 264.92 ± 68.04% (p<0.01) at concentrations of 20, 40, 60, 80, and 100 µg/mL, respectively, relative to the control. However, these changes did not follow a consistent trend across increasing concentrations of RE, as there was no clear dose-response relationship (Fig. 4a and 4b). RE demonstrated the most potent antiviral activity in cells treated after the 1 h infection period, as infectious virion production was significantly inhibited by every concentration tested (Fig. 5a, p<0.0001). There was a 70.90 ± 9.68% and 99.99 ± 0.01% reduction in viral titers in cells treated with 20 and 40 µg/mL of RE and a 100 ± 0.00% reduction in viral titers below the detectable limit in cells treated with 60, 80, and 100 µg/mL of RE (Fig. 5c, IC_50_ = 18.56 µg/mL, SI = 5.78). Genome copy numbers were also reduced by 95.87 ± 2.17% (p<0.0001), 99.60 ± 0.10% (p<0.0001), 99.61 ± 0.13% (p<0.0001), and 99.95 ± 0.03% (p<0.0001) at concentrations of 40, 60, 80, and 100 µg/mL, respectively (Fig. 5b and 5d, IC_50_ = 23.12 µg/mL, SI = 4.64). However, concentrations of 80 µg/mL and above were cytotoxic in HFF-1 cells treated with RE for 96 h (Fig. 1b). Lastly, pre-treatment of ZIKV with RE for 1 h prior to infection significantly increased viral titers by 65.99 ± 44.52% (p<0.05) at 20 µg/mL and reduced viral titers by 99.70 ± 0.30% (p<0.0001) at 60 µg/mL and 100 ± 0.00% (p<0.0001) at 80 and 100 µg/mL (Fig. 6a and 6c, IC_50_ = 56.25 µg/mL, SI = 2.14 µg/mL). Similarly, genome copy numbers decreased by 55.14 ± 22.42% (p<0.01), 99.84 ± 0.12% (p<0.0001), 99.90 ± 0.05% (p<0.0001), and 99.91 ± 0.06% (p<0.0001) at concentrations of 40, 60, 80, and 100 µg/mL, respectively (Fig. 6b and 6d, IC_50_ = 39.63 µg/mL, SI = 3.03 µg/mL). However, 100 µg/mL of RE was cytotoxic when HFF-1 cells were treated for 1 h followed by a 96 h incubation (Fig. 1a).

**Figure 3:**
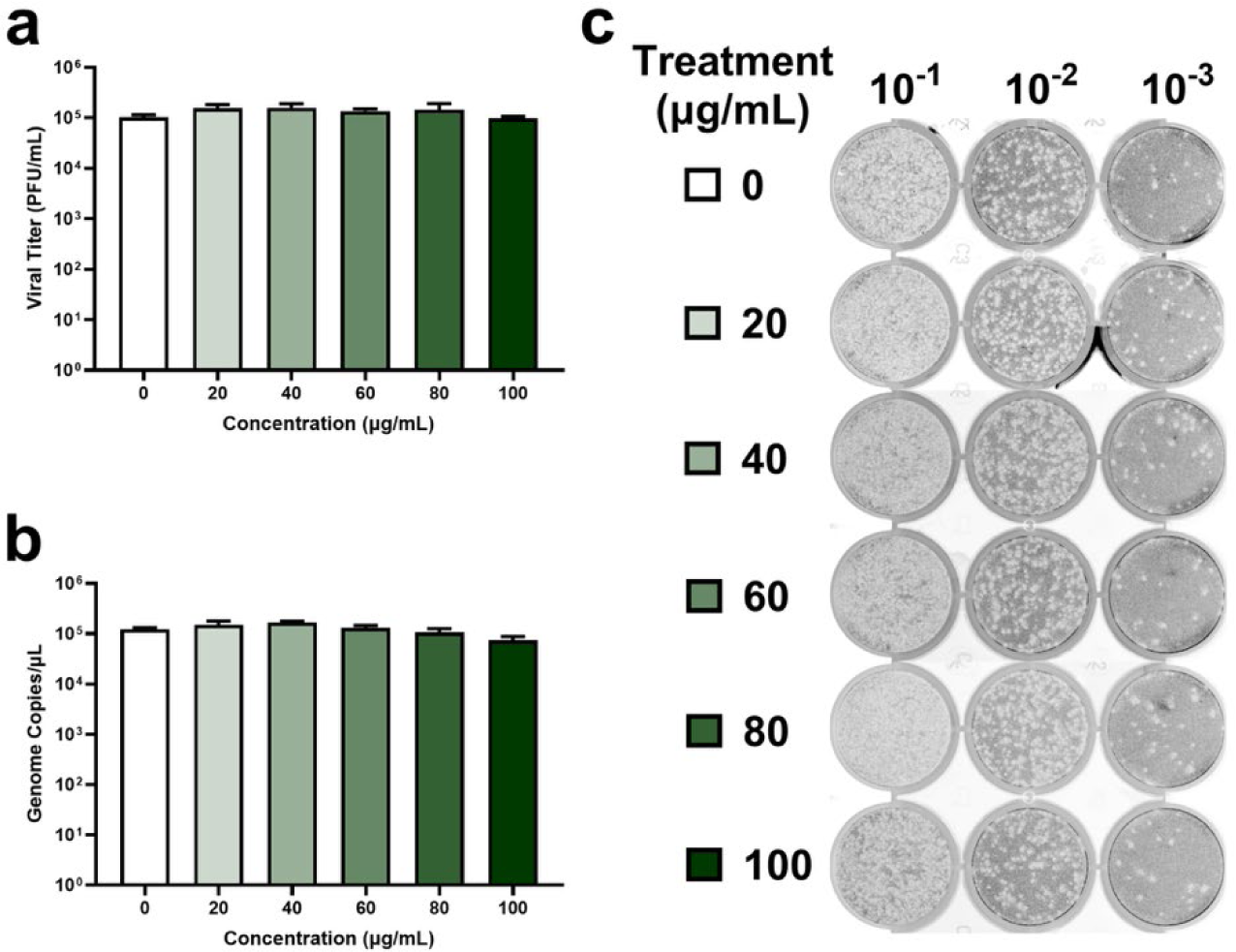
Treatment of HFF-1 cells with RE before infection does not impact ZIKV replication. (a) HFF-1 cells were treated with increasing concentrations of RE or a vehicle control (DMSO) for 1 h and subsequently infected with ZIKV (MOI = 1). Cell-free supernatants were collected 0 and 96 h post-infection, and the viral titer were quantified by plaque assay. Data are expressed as mean PFU/mL ± SEM for n = 3 independent experiments, each performed in duplicate. A two-way ANOVA and Dunnett’s multiple comparisons test were used to determine differences in viral titers following RE treatment relative to HFF-1 cells treated with DMSO (0 µg/mL). (b) Viral RNA was extracted from cell-free supernatants collected 96 h post-infection and quantified by RT-qPCR. Data are expressed as mean genome copies/mL ± SEM for n = 3 independent experiments, each performed in duplicate. A one-way ANOVA and Dunnett’s multiple comparisons test were used to determine differences in genome copy numbers following RE treatment relative to HFF-1 cells treated with DMSO (0 µg/mL). (c) Plaque assay images were taken using an Amersham Typhoon Biomolecular Imager and represent wells from one independent experiment in its entirety.

**Figure 4:**
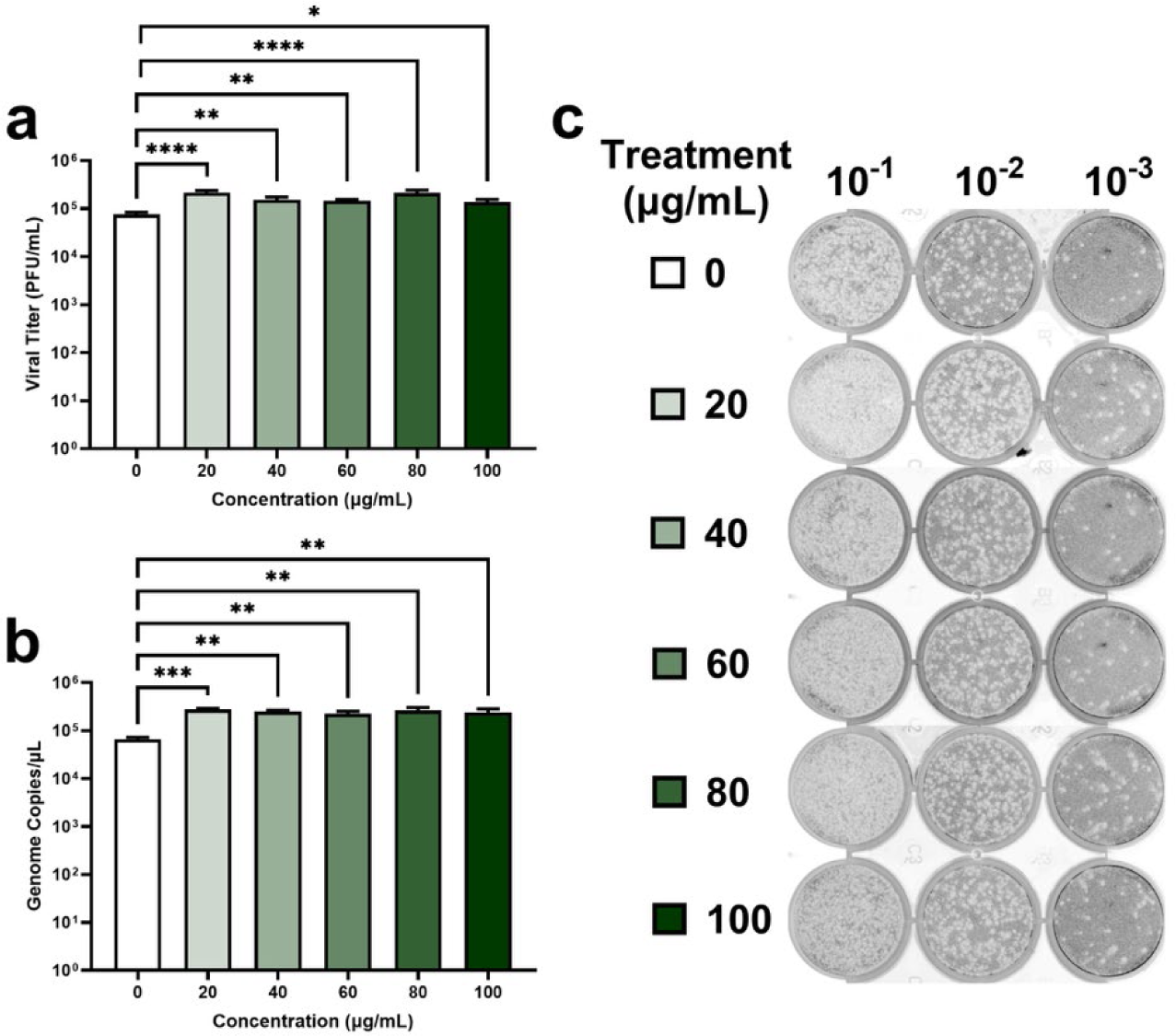
Treatment of HFF-1 cells with RE during infection enhances ZIKV replication. (a) HFF-1 cells were infected with ZIKV (MOI = 1) and simultaneously treated with increasing concentrations of RE or a vehicle control (DMSO) for 1 h. Cell-free supernatants were collected 0 and 96 h post-infection, and the viral titer of each sample was quantified by plaque assay. Data are expressed as mean PFU/mL ± SEM for n = 3 independent experiments, each performed in duplicate. A two-way ANOVA and Dunnett’s multiple comparisons test were used to determine differences in viral titers following RE treatment. *p<0.05, **p<0.01, ****p<0.0001 relative to HFF-1 cells treated with DMSO (0 µg/mL). (b) Viral RNA was extracted from cell-free supernatants collected 96 h post-infection and quantified by RT-qPCR. Data are expressed as mean genome copies/mL ± SEM for n = 3 independent experiments, each performed in duplicate. A one-way ANOVA and Dunnett’s multiple comparisons test were used to determine differences in genome copy numbers following RE treatment. **p<0.01, ***p<0.001 relative to HFF-1 cells treated with DMSO (0 µg/mL). (c) Plaque assay images were taken using an Amersham Typhoon Biomolecular Imager and represent wells from one independent experiment in its entirety.

**Figure 5:**
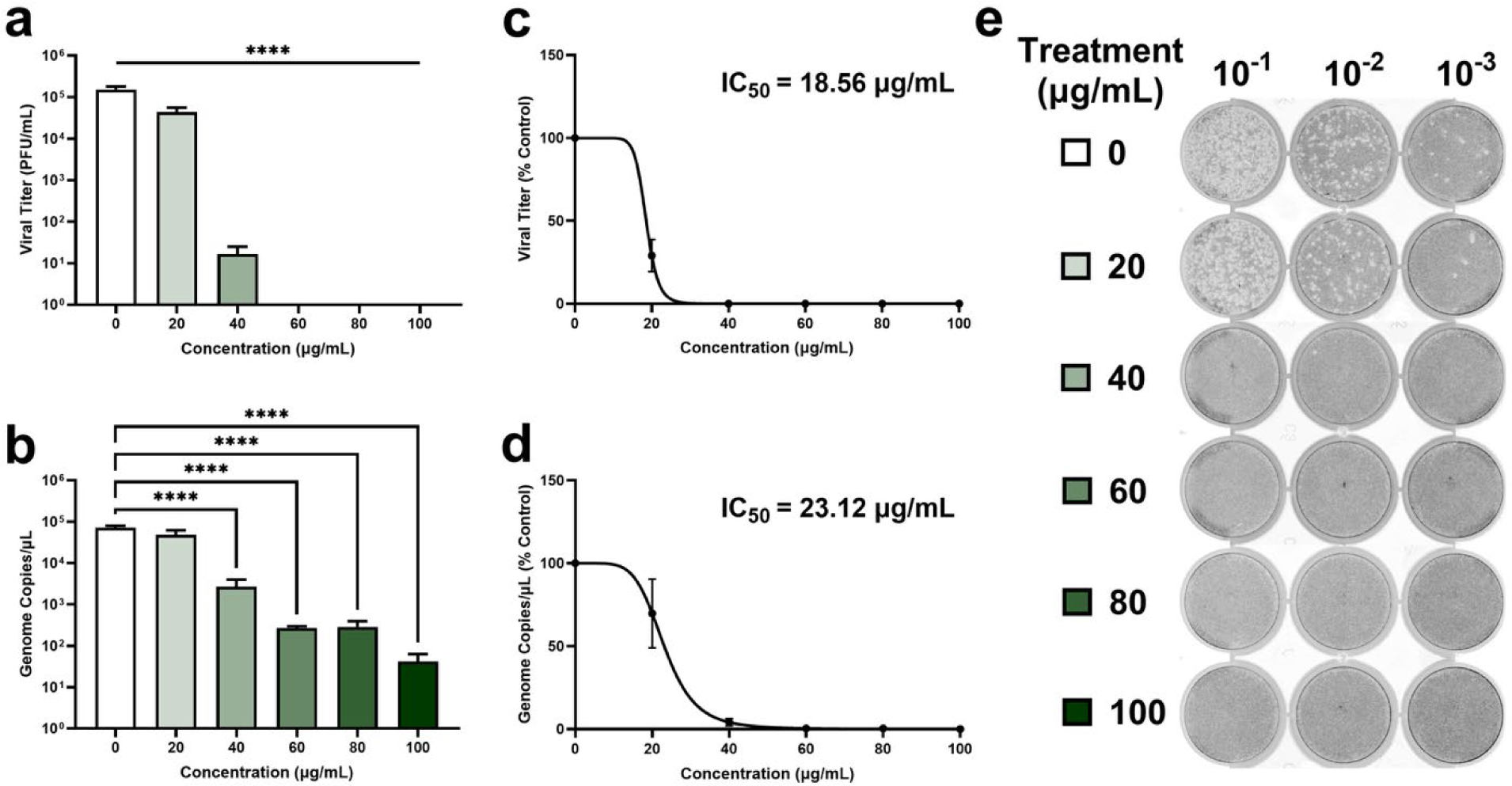
Treatment of HFF-1 cells with RE after infection inhibits ZIKV replication. (a) HFF-1 cells were infected with ZIKV (MOI = 1) for 1 h and subsequently treated with increasing concentrations of RE or a vehicle control (DMSO) for the entire post-infection period. Cell-free supernatants were collected 0 and 96 h post-infection and viral titers were quantified by plaque assay. Data are expressed as mean PFU/mL ± SEM for n = 3 independent experiments, each performed in duplicate. A two-way ANOVA and Dunnett’s multiple comparisons test were used to determine differences in viral titers following RE treatment. ****p<0.0001 relative to HFF-1 cells treated with DMSO (0 µg/mL). (b) Viral RNA was extracted from cell-free supernatants collected 96 h post-infection and quantified by RT-qPCR. Data are expressed as mean genome copies/mL ± SEM for n = 3 independent experiments, each performed in duplicate. A one-way ANOVA and Dunnett’s multiple comparisons test were used to determine differences in genome copy numbers following RE treatment. ****p<0.0001 relative to HFF-1 cells treated with DMSO (0 µg/mL). (c,d) IC_50_ values were calculated by non-linear regression analysis (inhibitor vs normalized response – variable slope). (e) Plaque assay images were taken using an Amersham Typhoon Biomolecular Imager and represent wells from one independent experiment in its entirety.

**Figure 6:**
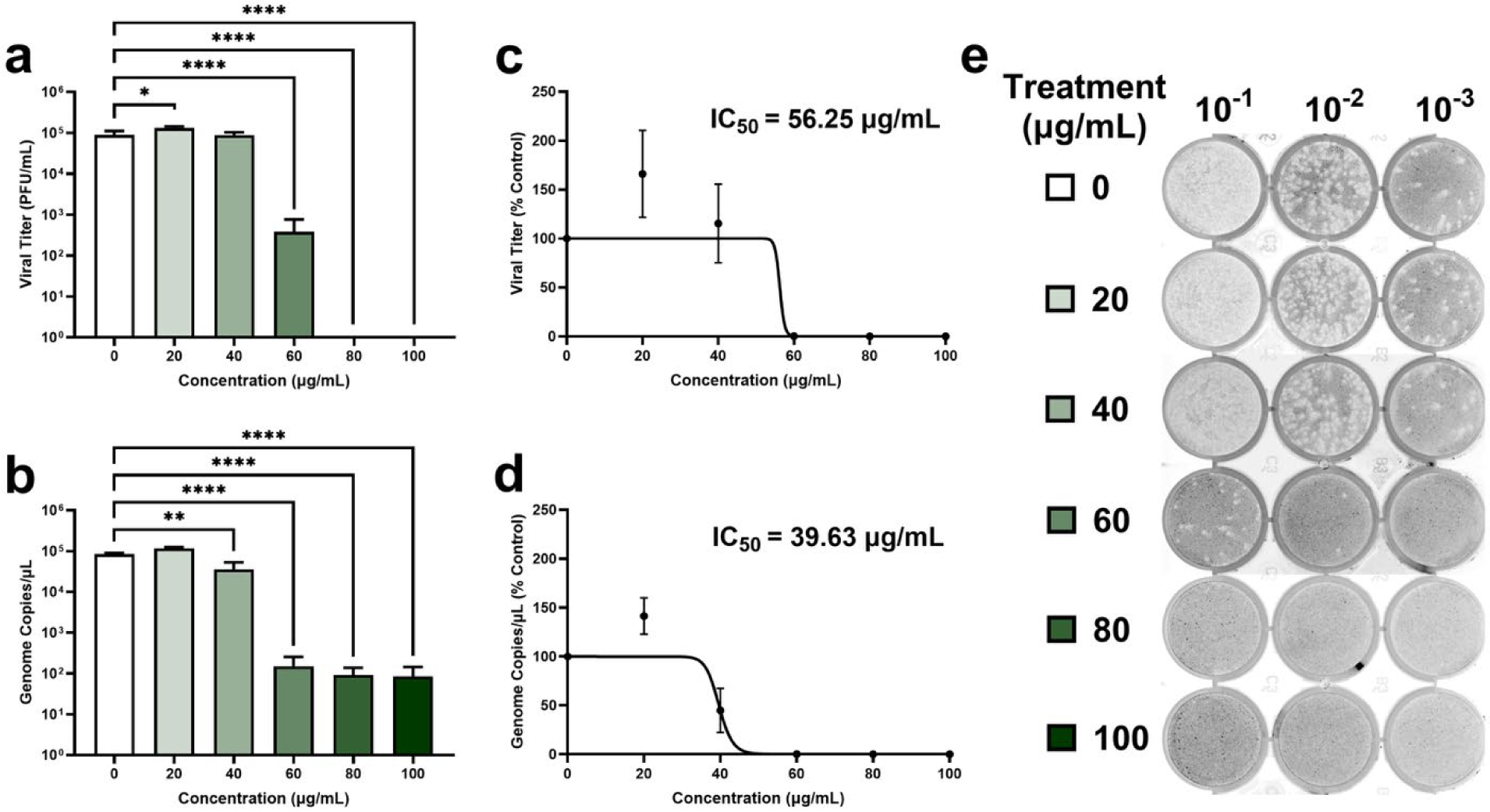
RE exhibits virucidal activity against ZIKV in HFF-1 cells. (a) ZIKV was incubated with increasing concentrations of RE or an equal volume of a vehicle control (DMSO) for 1 h. Subsequently, HFF-1 cells were infected with each sample (MOI = 1) and cell-free supernatants were collected 0 and 96 h post-infection. Viral titers were quantified by plaque assay. Data are expressed as mean PFU/mL ± SEM for n = 3 independent experiments, each performed in duplicate. A two-way ANOVA and Dunnett’s multiple comparisons test were used to determine differences in viral titers following RE treatment. *p<0.05, ****p<0.0001 relative to HFF-1 cells infected with DMSO treated ZIKV (0 µg/mL). (b) Viral RNA was extracted from cell-free supernatants collected 96 h post-infection and quantified by RT-qPCR. Data are expressed as mean genome copies/mL ± SEM for n = 3 independent experiments, each performed in duplicate. A one-way ANOVA and Dunnett’s multiple comparisons test were used to determine differences in genome copy numbers following RE treatment. **p<0.01, ****p<0.0001 relative to HFF-1 cells treated with DMSO (0 µg/mL). (c,d) IC_50_ values were calculated by non-linear regression analysis (inhibitor vs normalized response – variable slope). (e) Plaque assay images were taken using an Amersham Typhoon Biomolecular Imager and represent wells from one independent experiment in its entirety.

### CA and CO, but not RA, exhibit virucidal activity against ZIKV in Vero cells

To assess whether three main polyphenols in RE exert similar antiviral effects, ZIKV was pre-treated with 25 to 100 µM of each compound individually or an equivalent volume of DMSO for 1 h. The remaining infectivity of each sample was quantified by plaque assay. No effect on viral replication was observed when ZIKV was pre-treated with RA (Fig. 7a). However, there was a significant dose-dependent decrease in infectivity with CA (Fig. 8a) and CO (Fig. 9a). Treatment with CA resulted in a 30.69 ± 6.52% (p<0.01), 69.50 ± 5.23% (p<0.0001), and 99.27 ± 0.12% (p<0.0001) reduction in viral titers at concentrations of 25, 50, and 100 µM, respectively, corresponding to an IC_50_ of 34.83 µM (Fig. 8b). Similarly, CO caused a 67.04 ± 1.97% (p<0.001), 93.81 ± 1.64% (p<0.0001), and 98.96 ± 0.16% (p<0.0001) reduction in viral titers at concentrations of 25, 50, and 100 µM, respectively, with an IC_50_ of 19.55 µM (Fig. 9b).

**Figure 7:**
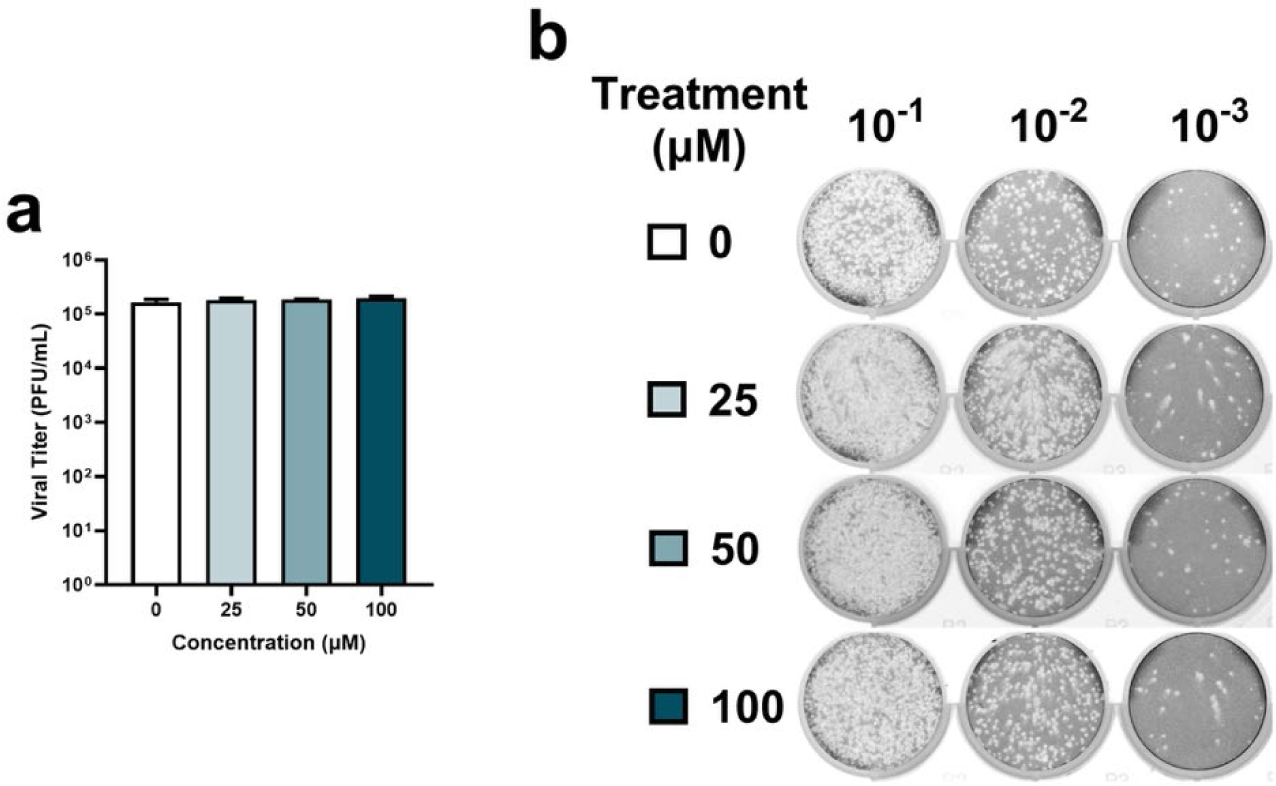
RA has no effect on ZIKV infection. (a) The virucidal activity of RA was evaluated by plaque reduction assay. ZIKV (290,000 PFU) was incubated with increasing concentrations of RA or an equal volume of a vehicle control (DMSO) for 1 h. Viral titers in each sample were then determined by plaque assay. Data are expressed as mean PFU/mL ± SEM for n = 3 independent experiments, each performed in duplicate. A one-way ANOVA and Dunnett’s multiple comparisons test were used to determine differences in viral titers following RA treatment relative to DMSO (0 µg/mL). (b) Plaque assay images were taken using an Amersham Typhoon Biomolecular Imager and represent wells from one independent experiment in its entirety.

**Figure 8:**
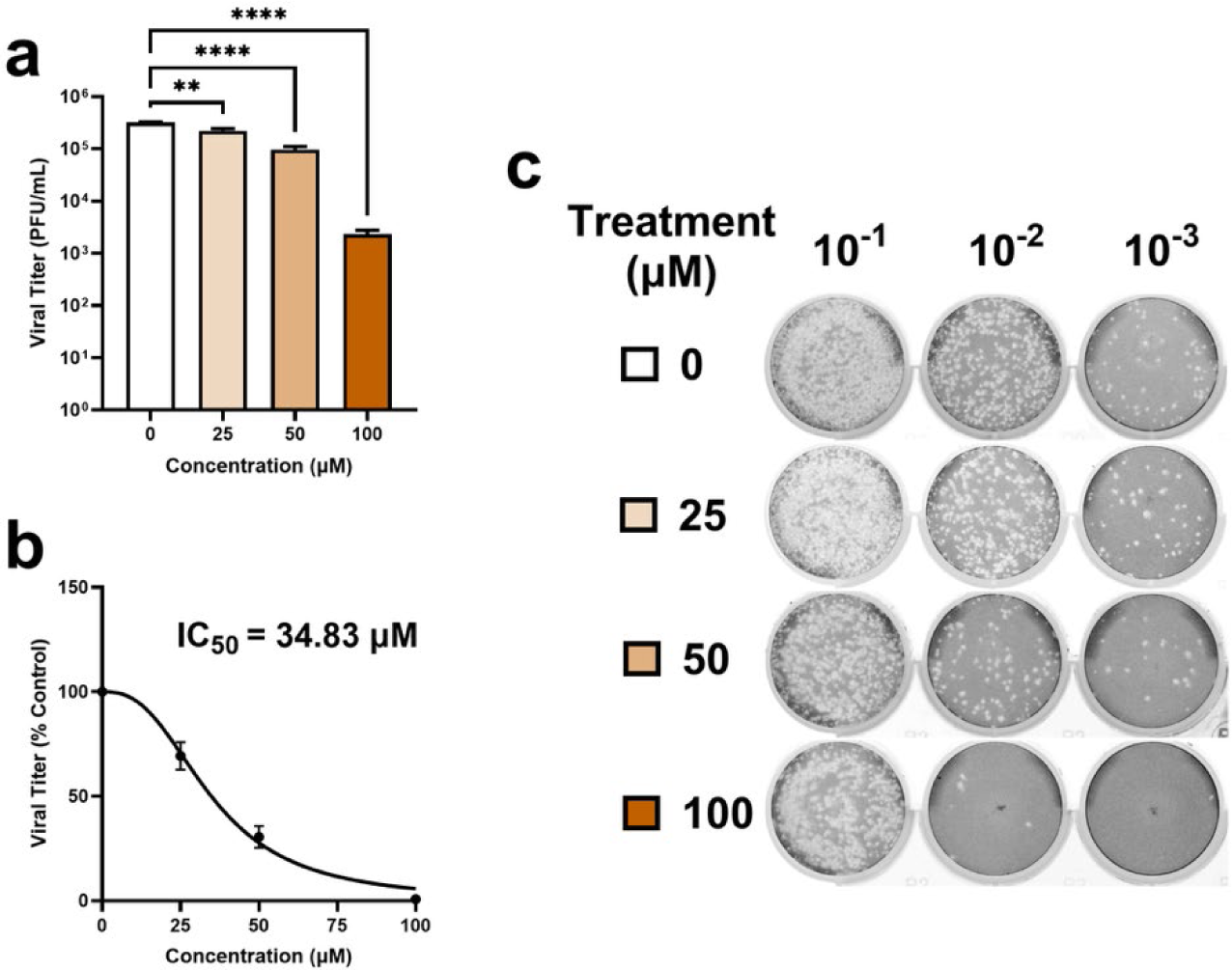
CA inhibits ZIKV infection in a dose-dependent manner. (a) The virucidal activity of CA was evaluated by plaque reduction assay. ZIKV (290,000 PFU) was incubated with increasing concentrations of CA or an equal volume of a vehicle control (DMSO) for 1 h. Viral titers in each sample were then determined by plaque assay. Data are expressed as mean PFU/mL ± SEM for n = 3 independent experiments, each performed in duplicate. A one-way ANOVA and Dunnett’s multiple comparisons test were used to determine differences in viral titers following CA treatment. **p<0.01, ****p<0.0001 relative to ZIKV incubated with DMSO (0 µg/mL). (b) The IC_50_ value was calculated by non-linear regression analysis (inhibitor vs normalized response – variable slope). (c) Plaque assay images were taken using an Amersham Typhoon Biomolecular Imager and represent wells from one independent experiment in its entirety.

**Figure 9:**
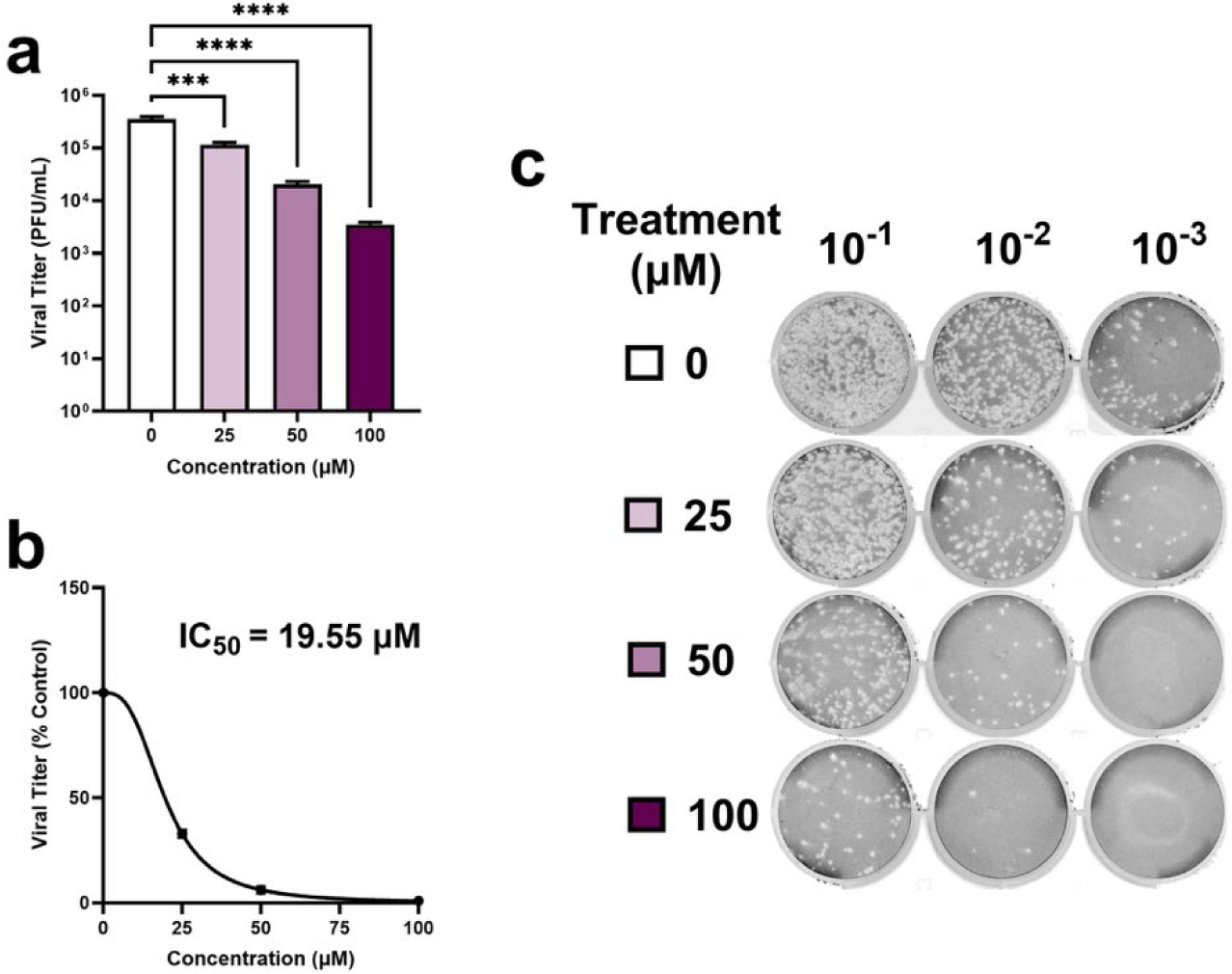
CO inhibits ZIKV infection in a dose-dependent manner. (a) The virucidal activity of CO was evaluated by plaque reduction assay. ZIKV (290,000 PFU) was incubated with increasing concentrations of CO or an equal volume of a vehicle control (DMSO) for 1 h. Viral titers in each sample were then determined by plaque assay. Data are expressed as mean PFU/mL ± SEM for n = 3 independent experiments, each performed in duplicate. A one-way ANOVA and Dunnett’s multiple comparisons test were used to determine differences in viral titers following CO treatment. ***p<0.001, ****p<0.0001 relative to ZIKV incubated with DMSO (0 µg/mL). (b) The IC_50_ value was calculated by non-linear regression analysis (inhibitor vs normalized response – variable slope). (c) Plaque assay images were taken using an Amersham Typhoon Biomolecular Imager and represent wells from one independent experiment in its entirety.

### HPLC analysis of RE

HPLC analysis was performed to identify and quantify CA, CO, and RA in the RE used in this study. A representative chromatogram for the identified compounds is shown in Fig. 10, and the corresponding results are presented in Table 1. The most abundant polyphenol per 1 mg of RE was CA (78.41 ± 4.64 µg/mg), followed by RA (27.69 ± 0.23 µg/mg) and CO (11.96 ± 0.32 µg/mg), with retention times of 17.62, 7.57, and 15.81 min, respectively. These amounts correspond to concentrations of 235.86 ± 13.96 µM CA, 76.85 ± 0.64 µM RA, and 36.20 ± 0.97 µM CO when 1 mg of RE is dissolved in 1 mL. Accordingly, at the lowest concentration of RE tested in the plaque reduction assay (25 µg/mL), ZIKV was exposed to approximately 5.90 ± 0.35 µM CA, 1.92 ± 0.02 µM RA, and 0.91 ± 0.02 µM CO.

**Figure 10:**
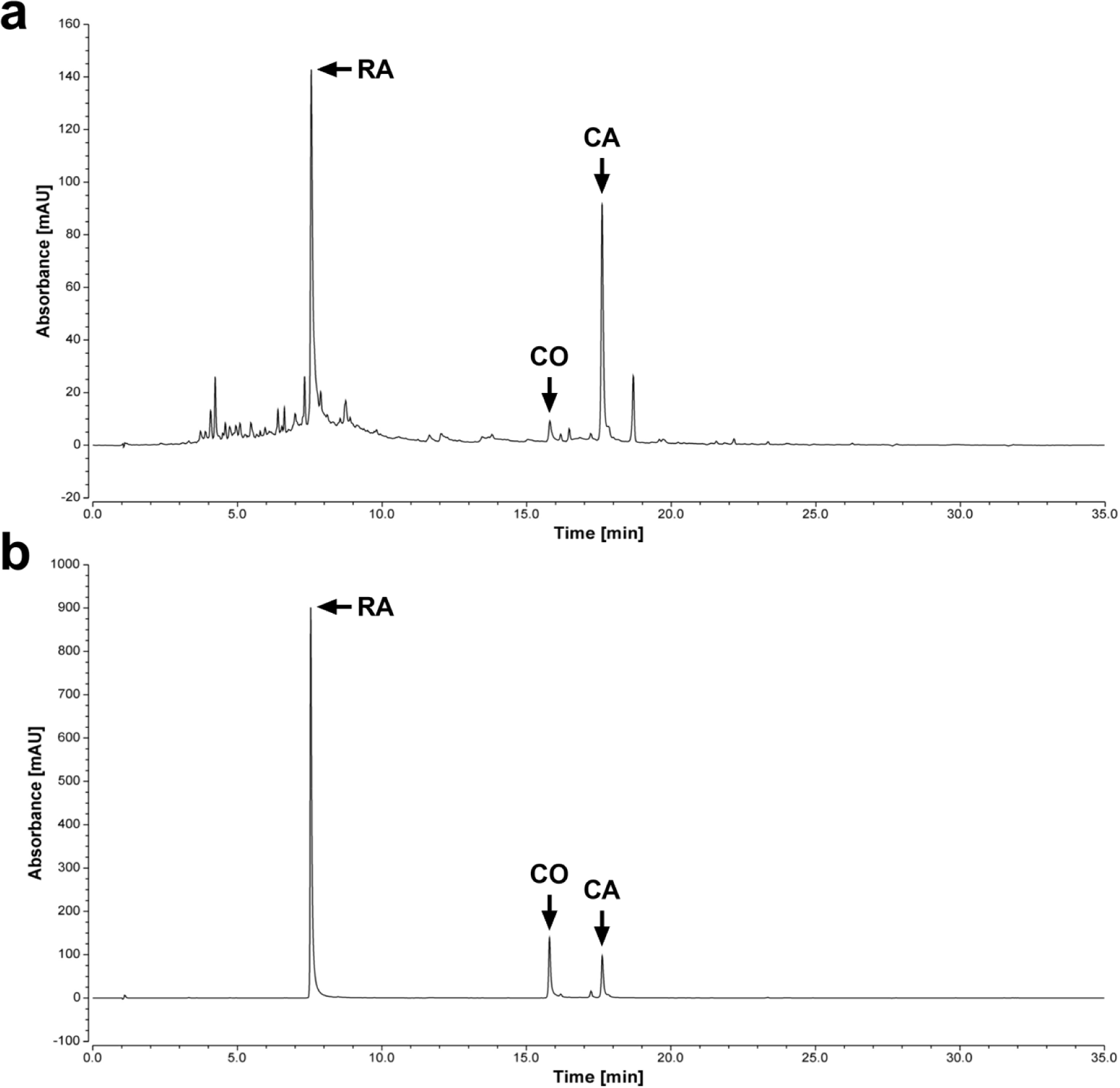
Separation and identification of major polyphenols in RE. Representative HPLC chromatograms of (a) RE and (b) a standard mixture of CA, CO, and RA. HPLC analysis was performed using a Kinetex C18 column and chromatograms were recorded at 284 nm. Peaks were identified by comparing retention times with authentic standards.

**Table 1:**
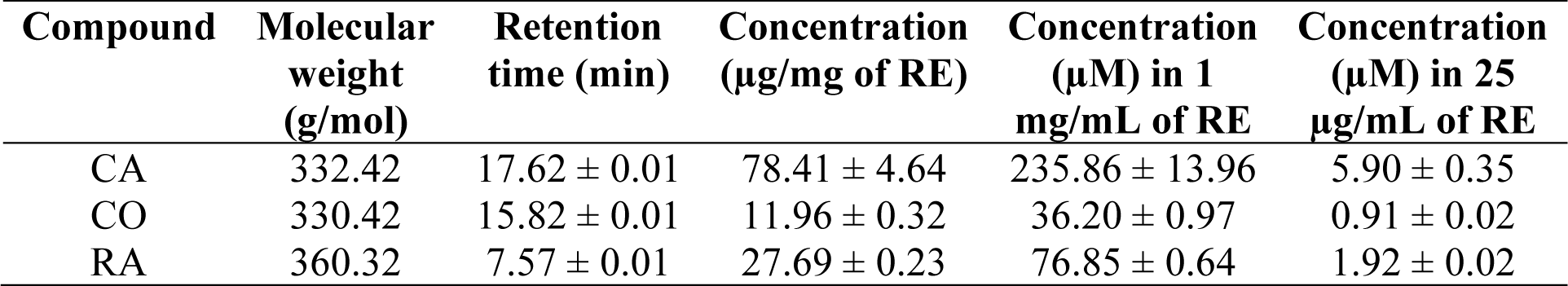
Quantification of major polyphenols in RE. HPLC analysis was performed using a Kinetex C18 column and detection at 284 nm. Concentrations of CA, CO, and RA were determined using the external standardization method with authentic standards. Data are expressed as mean ± SEM for n = 3 independent experiments.

| Compound | Molecular weight (g/mol) | Retention time (min) | Concentration ($\mu\text{g}/\text{mg}$ of RE) | Concentration ($\mu\text{M}$) in 1 mg/mL of RE | Concentration ($\mu\text{M}$) in 25 $\mu\text{g}/\text{mL}$ of RE |
| --- | --- | --- | --- | --- | --- |
| CA | 332.42 | $17.62 \pm 0.01$ | $78.41 \pm 4.64$ | $235.86 \pm 13.96$ | $5.90 \pm 0.35$ |
| CO | 330.42 | $15.82 \pm 0.01$ | $11.96 \pm 0.32$ | $36.20 \pm 0.97$ | $0.91 \pm 0.02$ |
| RA | 360.32 | $7.57 \pm 0.01$ | $27.69 \pm 0.23$ | $76.85 \pm 0.64$ | $1.92 \pm 0.02$ |

## Discussion

Although cases of ZIKV have significantly declined since reaching their peak in 2016, the virus continues to circulate at low levels in tropical and subtropical regions, posing an ongoing threat to human health^15^. Predictive models estimate that by 2050, 1.3 billion people may be at risk of ZIKV infection, as rising global temperatures continue to expand the distribution of competent vectors and create conditions suitable for ZIKV transmission^47^. Despite extensive efforts by the global scientific community, there are no vaccines or antiviral treatments against ZIKV currently available, leaving us unprepared for potential future outbreaks^3^. This study aimed to evaluate the antiviral potential of RE against ZIKV in human dermal fibroblasts, one of the earliest cellular targets of infection. Here, we provide novel evidence demonstrating RE exerts potent *in vitro* antiviral effects against ZIKV. RE was found to significantly impair ZIKV infection in Vero cells when the virus was treated before infection (Fig. 2). To assess the effects of RE on the ZIKV replication cycle, HFF-1 cells were treated before, during, and after infection or ZIKV was treated before infection. No antiviral activity was observed when HFF-1 cells were treated before (Fig. 3) or during infection (Fig. 4), however, ZIKV replication unexpectedly increased in the latter condition. There was a significant decrease in viral titers and genome copies when ZIKV was pre-treated with RE, indicating RE may target the initial stages of the viral replication cycle in HFF-1 cells (Fig. 6). RE was most effective when HFF-1 cells were treated post-infection, indicating RE may also target the later stages of the viral replication cycle (Fig. 5). Of the major compounds tested, CA (Fig. 8) and CO (Fig. 9), but not RA (Fig. 7), possess antiviral activity against ZIKV. To the best of our knowledge, this is the first study describing the antiviral activity of RE in the context of flavivirus infection, specifically ZIKV.

To determine the antiviral potential of RE against ZIKV, we employed a plaque reduction assay. This approach is commonly used in the literature^36,48–50^ to screen different compounds for antiviral activity and involves directly treating the virus before infecting a susceptible cell line, such as Vero. Here, we observed a significant dose-dependent decrease in viral replication with all concentrations of RE, with the lowest concentration causing a 52.77% reduction in viral titers (Fig. 2, IC_50_ = 25.06 µg/mL). Although very few studies have investigated the antiviral effects of RE, comparable results have been shown against HSV-1^30,32^ and HSV-2^30^ in Vero cells. However, Nolkemper et al. reported IC_50_ values of 0.646 µg/mL against HSV-1 and 1.055 µg/mL against HSV-2, which are much lower than those observed in the current study^30^. This discrepancy can be potentially attributed to differences between the viruses and procedures used. For example, Nolkemper et al. pre-treated 200 PFU of HSV before adding the entire inoculum to RC-37 cells (African green monkey kidney cells), whereas we pre-treated 290,000 PFU of ZIKV and infected Vero cells with serially diluted inoculum (10^-1^ to 10^-3^). Furthermore, variations in extraction solvents (water vs. methanol in our study) can substantially influence the chemical composition and corresponding bioactivity of RE^51^. Overall, our results are in general agreement with the literature, as several classes of natural products with anti-ZIKV activity exert a virucidal effect by directly interacting with virus particles or exhibit biological effects during viral adsorption and internalization^22^.

To identify which stage(s) of the ZIKV replication cycle are affected by RE, we performed four experiments using HFF-1 cells: 1) treatment of cells before infection, 2) simultaneous treatment and infection of cells, 3) treatment of cells after infection, and 4) treatment of ZIKV before infection. Fibroblasts are among the first cells that encounter ZIKV at the site of inoculation, therefore, HFF-1 cells were selected as a relevant infection model. Results show that treating cells before or during the adsorption phase does not impair ZIKV replication (Fig. 3 and 4). Nolkemper et al. found similar results, with no effect on HSV-2 and a minimal 24% reduction in HSV-1 replication when RC-37 cells were treated with a non-cytotoxic concentration of RE before infection. However, treatment of RC-37 cells during infection caused a 36% and 67% reduction in HSV-1 and HSV-2 replication, respectively^30^. In another study, 20 µg/mL RE caused a 66-fold reduction in hRSV production in A549 cells treated before infection, conflicting with our results, though a different virus and cell type^29^. Moreover, Lin *et al.* demonstrated a significant 21.9% reduction in viral VP1 mRNA and an 88.6% reduction in EV71 replication in human rhabdomyosarcoma (RD) cells treated with 10 µM of RA, one of the main constituents found in RE, during infection. Similar to our findings, RA minimally inhibited EV71 by 5.54% in RD cells treated before infection^52^. Surprisingly, we observed a statistically significant increase in ZIKV replication in cells treated during infection with every concentration of RE (Fig. 4), as well as in cells infected with ZIKV pre-treated with 20 µg/mL (Fig. 5). As far as we know, this effect has not been reported in the literature. Considering that pH can have a significant impact on ZIKV infectivity^53^, we measured the pH of various concentrations of RE prepared in DMEM. No major changes in pH were observed in comparison to DMSO, with all values falling within a range of approximately 8.2 to 8.5 (data not shown). Therefore, the increase in virus production may have occurred due to an experimental artefact, but further investigation is required to determine if underlying mechanisms are at play.

Unlike the results above, RE was shown to exert potent antiviral activity when cells were treated post-infection. There was a considerable 70.90% decrease in infectious virion production (IC_50_ = 18.56 µg/mL, SI = 5.78) at the lowest concentration tested (20 µg/mL) and a 95.87% decrease in genome copy numbers (IC_50_ = 23.12 µg/mL, SI = 4.64) at 40 µg/mL, with viral titers falling below the detectable limit at 60 µg/mL and beyond (Fig. 5). This is in agreement with Horvath et al., who found a 90.46% reduction in HSV-2 genome concentrations in Vero cells treated with 0.47 mg/mL of RE after infection^33^. Similarly, Al-Megrin *et al.* reported that 30 and 40 µg/mL of RE reduces plaque formation of HSV-1 by 55% and HSV-2 by 65%, respectively, when incorporated into the overlay^32^. In contrast to these findings, RE had no effect on HSV-1 or HSV-2 plaque formation in RC-37 cells^30^. Of the main polyphenols found in RE, CA has been shown to reduce HSV-2^33^ and hRSV^29^ genome concentrations in cells treated post-infection, whereas EV71^52^ mRNA and progeny virus production were reduced in RA treated cells, further supporting our results. Based on an *in-silico* study, RE may interfere with the NS5 RNA-dependent RNA polymerase (RdRp), which is essential for replicating the ZIKV genome. This is due to inhibitory potential of CA (-6.8 kcal/mol), CO (-7.0 kcal/mol), RA (6.9 kcal/mol), and several other compounds, which were predicted to have high binding energy against the RdRp, although *in vitro* studies are required to validate these data^26^. Collectively, these data suggest that RE may act on the later stages of the viral replication cycle, however, further research is necessary to determine the underlying mechanisms, such as effects on gene expression or virion assembly and release.

As seen with the plaque reduction assay, pre-treatment of ZIKV with RE significantly inhibited infection in HFF-1 cells, resulting in a 99.70% reduction in progeny virions at a concentration of 60 µg/mL and complete inhibition at 80 µg/mL (Fig. 6, IC_50_ = 56.25 µg/mL, SI = 2.14). Viral RNA was also significantly reduced by 55.14%, 99.84%, 99.90%, and 99.91% at concentrations of 40, 60, 80, and 100 µg/mL, respectively (IC_50_ = 39.63 µg/mL, SI = 3.03 µg/mL). Similar results were reported for HSV pre-treated with the maximum non-cytotoxic concentration of RE (exact concentration not specified) in RC-37 cells, resulting in 97% and 99% reductions in HSV-1 and HSV-2, respectively^30^. Unlike our plaque reduction assay results, no decrease in viral titers was observed when ZIKV was pre-treated with concentrations below 60 µg/mL, which may be due to the distinct species and tissue origin of each cell line, as well as fundamental differences between the assays. For example, ZIKV was treated with RE for 1 h and the infectivity of each sample was quantified by plaque reduction assay, whereas in the time-of-addition analysis, plaque assay was used to quantify the release of infectious virions by HFF-1 cells after infection with pre-treated ZIKV. Since no antiviral effects were observed in HFF-1 cells treated during infection, these findings indicate that RE directly interacts with extracellular ZIKV particles by potentially binding to viral proteins involved in mediating viral entry or damaging virions, thereby impairing their ability to infect host cells.

To determine whether three major polyphenols in RE exert anti-ZIKV activity, each compound was individually screened using a plaque reduction assay. Pre-treatment of ZIKV with CA and CO significantly reduced ZIKV plaque formation in a dose-dependent manner, with IC_50_ values of 34.83 and 19.55 µM, respectively (Fig. 8 and 9). Although both compounds are phenolic diterpenoids bearing a catechol moiety, CA carries a carboxyl moiety, whereas CO carries a lactone moiety^54^. These structural differences suggest that the lactone moiety may contribute to the lower IC_50_ of CO. In comparison, RA had no effect on ZIKV infection at any concentration tested (Fig. 7). This was somewhat unexpected, as RA is the only compound with existing evidence of antiviral activity against flaviviruses. Pre-treatment of DENV with RA has been shown to inhibit all four serotypes *in vitro*^36^, as well as significantly reduce mortality, viral mRNA, and proinflammatory cytokine levels in mice peritoneally treated with RA and infected with a lethal dose of JEV^37^. There is also *in vitro* evidence demonstrating the efficacy of RA against the influenza A virus H1N1^48^, EV71^52,55^, and hepatitis B virus^56^, as well as *in vivo* evidence against a lethal dose of EV71 in mice^57^. Based on the structural similarities among flaviviruses, we anticipated that RA would exert antiviral activity against ZIKV. Its failure to do so indicates that its effects may be specific to DENV and JEV. The antiviral activity of CA has also been well characterized, as it was shown to be effective against HSV-2^33^, hRSV^29^, and influenza A virus H1N1^48^. However, another study reported no effect against influenza A virus H1N1^29^. Interestingly, we did not anticipate CO to have such a profound effect, as only two groups have analyzed its antiviral properties with conflicting results^29,58^. HPLC analysis revealed that the lowest concentration of RE tested by plaque reduction assay (25 µg/mL), which caused a 52.77% reduction in viral titers (Fig. 2), contained approximately 5.90 µM CA and 0.91 µM CO (Table 1). These concentrations are substantially lower than the IC_50_ values determined for each compound individually, suggesting that additional compounds within RE, or synergistic interactions between compounds, underlie its antiviral activity and warrant further investigation.

There are several limitations worth considering. Many antiviral studies include a positive control to validate assay performance and ensure the proper interpretation of results. Since no clinically approved antiviral exists for ZIKV, we did not include a positive control, although a repurposed drug with *in vitro* anti-ZIKV activity could theoretically serve this purpose. Secondly, depending on soil characteristics, climate, light exposure, and other growing conditions, the chemical composition of plant extracts may vary, limiting their reproducibility. To address this, studies should report the composition of each extract, where possible, to facilitate comparisons across the literature. Lastly, while *in vitro* studies provide valuable insights and are necessary during the early stages of investigation, the observed effects may not directly translate to humans. The human body consists of many diverse cell types interacting within complex tissues and organ systems, and antiviral efficacy can be influenced by metabolism, immune responses, bioavailability, and other factors. Therefore, it is important to evaluate antivirals in small animal models, as these are more physiologically relevant. However, a lack of immunocompetent animal models for ZIKV and other flaviviruses remains a challenge that must be addressed moving forward.

As mosquitoes and flaviviruses continue to expand into new regions and increasingly threaten human health, new antiviral agents are urgently needed. In this study, we demonstrate that RE exhibits potent antiviral activity against ZIKV by directly interacting with virus particles before infection in Vero and HFF-1 cells and interfering with post-entry processes of the ZIKV replication cycle in HFF-1 cells. Screening three major polyphenols found in RE revealed that CA and CO significantly reduce ZIKV infectivity and are promising candidates for follow-up studies. Whether RE can also inhibit other medically important flaviviruses, such as DENV and WNV, remains to be determined. Collectively, these findings provide novel evidence supporting the antiviral potential of RE and its individual components, warranting further investigation into their mechanisms of action and effects in additional cell and viral models.

## Authorship Contributions

JTM, RWEC, JMC, EIP, and AJM conceived and designed the study. RWEC, JMC, and DD assisted with experimental planning. JTM performed the experiments with assistance from RWEC, NJH, and CAL. JTM and AJM analyzed the data and wrote the manuscript. All authors contributed to discussion and editing of the manuscript.

## Abbreviations

CC_50_: 50% cytotoxic concentration
CA: Carnosic acid
CMC: Carboxymethylcellulose
CO: Carnosol
DENV: Dengue virus
DMEM: Dulbecco’s modified Eagle’s medium
DMSO: Dimethyl sulfoxide
EV71: Enterovirus 71
HFF-1: Human foreskin fibroblast cells
hRSV: Human respiratory syncytial virus
HSV: Herpes simplex virus
IC_50_: 50% inhibitory concentration
JEV: Japanese encephalitis virus
MOI: Multiplicity of infection
NS: Non-structural
RT-qPCR: Reverse transcription-quantitative polymerase chain reaction
PBS: Phosphate-buffered saline
Pen/strep: penicillin/streptomycin
PFU: Plaque forming unit
RA: Rosmarinic acid
RD: Human rhabdomyosarcoma
RdRp: RNA-dependent RNA polymerase
RE: Rosemary extract
SI: Selectivity index
WHO: World Health Organization
WNV: West Nile virus
YFV: Yellow fever virus
ZIKV: Zika virus

---

**Supplemental Figure 1:**
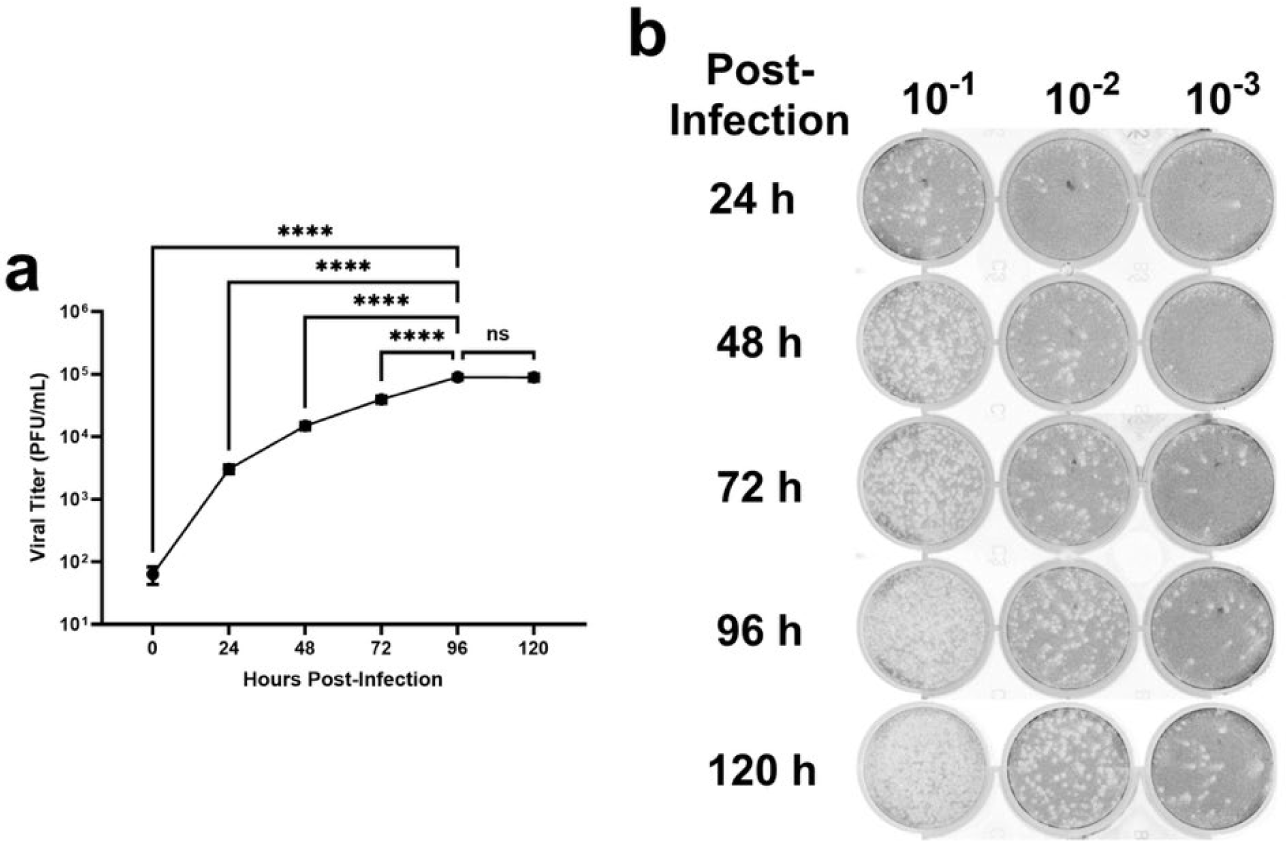
Infectious titers peak at 96 h in HFF-1 cells infected with ZIKV. (a) HFF-1 cells were infected with ZIKV (MOI = 1) for 1 h at 37°C. Cell-free supernatants were collected 0, 24, 48, 72, 96, and 120 h post infection and viral titers in each sample were quantified by plaque assay. Data are expressed as PFU/mL ± SEM for n = 3 independent experiments, each performed in duplicate. A two-way ANOVA and Dunnett’s multiple comparisons test were used to determine differences in viral titers between each timepoint. ****p<0.0001 relative to supernatant collected 96 h post-infection. (b) Plaque assay images were taken using an Amersham Typhoon Biomolecular Imager and represent wells from one independent experiment in its entirety.

## References

1. Vázquez-Calvo, Á., Jiménez de Oya, N., Martín-Acebes, M. A., Garcia-Moruno, E. & Saiz, J.-C. Antiviral Properties of the Natural Polyphenols Delphinidin and Epigallocatechin Gallate against the Flaviviruses West Nile Virus, Zika Virus, and Dengue Virus. Front. Microbiol. 8, (2017).

2. Noorbakhsh, F. et al. Zika Virus Infection, Basic and Clinical Aspects: A Review Article. Iran. J. Public Health 48, 20–31 (2019).

3. Pielnaa, P. et al. Zika virus-spread, epidemiology, genome, transmission cycle, clinical manifestation, associated challenges, vaccine and antiviral drug development. Virology 543, 34–42 (2020).

4. Duffy, M. R. et al. Zika Virus Outbreak on Yap Island, Federated States of Micronesia. N. Engl. J. Med. 360, 2536–2543 (2009).

5. Zhao, R. et al. Flavivirus: From Structure to Therapeutics Development. Life 11, 615 (2021).

6. Laureti, M., Narayanan, D., Fazakerley, J. K. & Kedzierski, L. Flavivirus Receptors: Diversity, Identity, and Cell Entry. Front. Immunol. 9, (2018).

7. Gorshkov, K. et al. Zika Virus: Origins, Pathological Action, and Treatment Strategies. Front. Microbiol. 9, 3252 (2019).

8. Bustos-Arriaga, J. et al. Soluble mediators produced by the crosstalk between microvascular endothelial cells and dengue-infected primary dermal fibroblasts inhibit dengue virus replication and increase leukocyte transmigration. Immunol. Res. 64, 392–403 (2016).

9. Garcia, M., Wehbe, M., Lévêque, N. & Bodet, C. Skin innate immune response to flaviviral infection. Eur. Cytokine Netw. 28, 41–51 (2017).

10. Hamel, R. et al. Biology of Zika Virus Infection in Human Skin Cells. J. Virol. 89, 8880– 8896 (2015).

11. Dick, M. K., Miao, J. H. & Limaiem, F. Histology, Fibroblast. StatPearls (2024).

12. Zika epidemiology update - May 2024. https://www.who.int/publications/m/item/zika-epidemiology-update-may-2024 (2024).

13. Komarasamy, T. V., Adnan, N. A. A., James, W. & Balasubramaniam, V. R. Finding a chink in the armor: Update, limitations, and challenges toward successful antivirals against flaviviruses. PLoS Negl. Trop. Dis. 16, e0010291 (2022).

14. Sadeer, N. B. et al. Secondary metabolites as potential drug candidates against Zika virus, an emerging looming human threat: Current landscape, molecular mechanism and challenges ahead. J. Infect. Public Health 16, 754–770 (2023).

15. Musso Didier, Ko Albert I., & Baud David. Zika Virus Infection — After the Pandemic. N. Engl. J. Med. 381, 1444–1457 (2019).

16. Grigalunas, M., Brakmann, S. & Waldmann, H. Chemical Evolution of Natural Product Structure. J. Am. Chem. Soc. 144, 3314–3329 (2022).

17. Annunziata, G. et al. May Polyphenols Have a Role Against Coronavirus Infection? An Overview of in vitro Evidence. Front. Med. 7, (2020).

18. Dzobo, K. The Role of Natural Products as Sources of Therapeutic Agents for Innovative Drug Discovery. Compr. Pharmacol. 408–422 (2022).

19. Patridge, E., Gareiss, P., Kinch, M. S. & Hoyer, D. An analysis of FDA-approved drugs: natural products and their derivatives. Drug Discov. Today 21, 204–207 (2016).

20. Dias, D. A., Urban, S. & Roessner, U. A Historical Overview of Natural Products in Drug Discovery. Metabolites 2, 303–336 (2012).

21. Zhou, B. & Yue, J.-M. Natural products are the treasure pool for antimalarial agents. Natl. Sci. Rev. 9, nwac112 (2022).

22. Pereira, R. S. et al. Natural Products and Derivatives as Potential Zika virus Inhibitors: A Comprehensive Review. Viruses 15, 1211 (2023).

23. Lee, J. L., Loe, M. W. C., Lee, R. C. H. & Chu, J. J. H. Antiviral activity of pinocembrin against Zika virus replication. Antiviral Res. 167, 13–24 (2019).

24. Shiravi, A. et al. Rosemary and its protective potencies against COVID-19 and other cytokine storm associated infections: A molecular review. Mediterr. J. Nutr. Metab. 14, 401– 416 (2021).

25. Harlina, P. W., Ma, M., Shahzad, R. & Khalifa, I. Effect of Rosemary Extract on Lipid Oxidation, Fatty Acid Composition, Antioxidant Capacity, and Volatile Compounds of Salted Duck Eggs. Food Sci. Anim. Resour. 42, 689–711 (2022).

26. Activity of Phytochemical Constituents of Black Pepper and Rosemary Against Zika Virus: An In silico Approach. Lett. Appl. NanoBioScience 11, 3393–3404 (2021).

27. Zhang, L. & Lu, J. Rosemary (*Rosmarinus officinalis* L.) polyphenols and inflammatory bowel diseases: Major phytochemicals, functional properties, and health effects. Fitoterapia 177, 106074 (2024).

28. Nieto, G., Ros, G. & Castillo, J. Antioxidant and Antimicrobial Properties of Rosemary (Rosmarinus officinalis, L.): A Review. Medicines 5, 98 (2018).

29. Shin, H.-B. et al. Antiviral activity of carnosic acid against respiratory syncytial virus. Virol. J. 10, 303 (2013).

30. Nolkemper, S., Reichling, J., Stintzing, F. C., Carle, R. & Schnitzler, P. Antiviral Effect of Aqueous Extracts from Species of the Lamiaceae Family against Herpes simplex Virus Type 1 and Type 2 in vitro. Planta Med. 72, 1378–1382 (2006).

31. Mancini, D. A. P., Torres, R. P., Pinto, J. R. & Mancini-Filho, J. Inhibition of DNA virus: Herpes-1 (HSV-1) in cellular culture replication, through an antioxidant treatment extracted from rosemary spice. *Braz*. J. Pharm. Sci. 45, 127–133 (2009).

32. AL-Megrin, W. A., et al. Potential antiviral agents of Rosmarinus officinalis extract against herpes viruses 1 and 2. Biosci. Rep. 40, BSR20200992 (2020).

33. Horváth, G. et al. Carnosic Acid Inhibits Herpes Simplex Virus Replication by Suppressing Cellular ATP Synthesis. Int. J. Mol. Sci. 25, 4983 (2024).

34. Ahmed, S. R., Banik, A., Anni, S. M. & Chowdhury, M. M. H. Inhibitory potential of plant-derived metabolites against Zika virus: a computational-aided approach. Phytomedicine Plus 1, 100129 (2021).

35. Samy, C. R. A., et al. (R)-(+)-Rosmarinic Acid as an Inhibitor of Herpes and Dengue Virus Replication: an In Silico Assessment. Rev. Bras. Farmacogn. 33, 543–550 (2023).

36. Panchal, R. et al. Antiviral Activity of Rosmarinic Acid Against Four Serotypes of Dengue Virus. Curr. Microbiol. 79, 203 (2022).

37. Swarup, V., Ghosh, J., Ghosh, S., Saxena, A. & Basu, A. Antiviral and Anti-Inflammatory Effects of Rosmarinic Acid in an Experimental Murine Model of Japanese Encephalitis. Antimicrob. Agents Chemother. 51, 3367–3370 (2007).

38. Mena, P. et al. Phytochemical Profiling of Flavonoids, Phenolic Acids, Terpenoids, and Volatile Fraction of a Rosemary (Rosmarinus officinalis L.) Extract. Molecules 21, 1576 (2016).

39. Borrás-Linares, I. et al. Rosmarinus Officinalis Leaves as a Natural Source of Bioactive Compounds. Int. J. Mol. Sci. 15, 20585–20606 (2014).

40. Kheiria, H. et al. Total Phenolic Content and Polyphenolic Profile of Tunisian Rosemary (*Rosmarinus officinalis* L.) Residues. Natural Drugs from Plants (2021).

41. Cedeño-Pinos, C., Martínez-Tomé, M., Murcia, M. A., Jordán, M. J. & Bañón, S. Assessment of Rosemary (Rosmarinus officinalis L.) Extract as Antioxidant in Jelly Candies Made with Fructan Fibres and Stevia. Antioxidants 9, 1289 (2020).

42. Coish, J. M. et al. Zika Virus Replication in a Mast Cell Model is Augmented by Dengue Virus Antibody-Dependent Enhancement and Features a Selective Immune Mediator Secretory Profile. Microbiol. Spectr. 10, e01772-22 (2022).

43. Gaudry, A. et al. The Flavonoid Isoquercitrin Precludes Initiation of Zika Virus Infection in Human Cells. Int. J. Mol. Sci. 19, 1093 (2018).

44. Cirne-Santos, C. C. et al. In vitro antiviral activity against Zika virus from a natural product of the Brazilian red seaweed Bryothamnion triquetrum. Acta Virol. 65, 402–410 (2022).

45. Saivish, M. V. et al. Antiviral Activity of Quercetin Hydrate against Zika Virus. Int. J. Mol. Sci. 24, 7504 (2023).

46. Vázquez-Calvo, Á., Jiménez de Oya, N., Martín-Acebes, M. A., Garcia-Moruno, E. & Saiz, J.-C. Antiviral Properties of the Natural Polyphenols Delphinidin and Epigallocatechin Gallate against the Flaviviruses West Nile Virus, Zika Virus, and Dengue Virus. Front. Microbiol. 8, 1314 (2017).

47. Ryan, S. J. et al. Warming temperatures could expose more than 1.3 billion new people to Zika virus risk by 2050. Glob. Change Biol. 27, 84–93 (2021).

48. Elebeedy, D., et al. *In vitro* and computational insights revealing the potential inhibitory effect of Tanshinone IIA against influenza A virus. Comput. Biol. Med. 141, 105149 (2022).

49. Koch, C., Reichling, J., Schneele, J. & Schnitzler, P. Inhibitory effect of essential oils against herpes simplex virus type 2. Phytomedicine 15, 71–78 (2008).

50. Vázquez-Calvo, Á., Jiménez de Oya, N., Martín-Acebes, M. A., Garcia-Moruno, E. & Saiz, J.-C. Antiviral Properties of the Natural Polyphenols Delphinidin and Epigallocatechin Gallate against the Flaviviruses West Nile Virus, Zika Virus, and Dengue Virus. Front. Microbiol. 8, (2017).

51. Al-jaafreh, A. M. Evaluation of Antioxidant Activities of Rosemary (Rosmarinus officinalis L.) Essential Oil and Different Types of Solvent Extractions. Biomed. Pharmacol. J. 17, 323– 339 (2024).

52. Lin, W.-Y., Yu, Y.-J. & Jinn, T.-R. Evaluation of the virucidal effects of rosmarinic acid against enterovirus 71 infection via in vitro and in vivo study. Virol. J. 16, 94 (2019).

53. Müller, J. A. et al. Inactivation and Environmental Stability of Zika Virus. Emerg. Infect. Dis. 22, 1685–1687 (2016).

54. Mantzourani, C., Tarantilis, P. A. & Kokotou, M. G. Carnosic Acid and Carnosol: Analytical Methods for Their Determination in Plants, Foods and Biological Samples. Separations 10, 481 (2023).

55. Chung, Y.-C. et al. Magnesium lithospermate B and rosmarinic acid, two compounds present in *Salvia miltiorrhiza*, have potent antiviral activity against enterovirus 71 infections. Eur. J. Pharmacol. 755, 127–133 (2015).

56. Tsukamoto, Y. et al. Rosmarinic acid is a novel inhibitor for Hepatitis B virus replication targeting viral epsilon RNA-polymerase interaction. PLOS ONE 13, e0197664 (2018).

57. Hsieh, C.-F. et al. Rosmarinic acid exhibits broad anti-enterovirus A71 activity by inhibiting the interaction between the five-fold axis of capsid VP1 and cognate sulfated receptors. Emerg. Microbes Infect. 9, 1194–1205.

58. Aruoma, O. I. et al. An evaluation of the antioxidant and antiviral action of extracts of rosemary and provençal herbs. Food Chem. Toxicol. 34, 449–456 (1996).

